# Acquired and intrinsic resistance to vemurafenib in BRAF^V600E^-driven melanoma brain metastases

**DOI:** 10.1101/2022.12.20.521202

**Authors:** Ping Zhang, Laura E. Kuil, Levi C.M. Buil, Stephan Freriks, Jos H. Beijnen, Olaf van Tellingen, Mark C. de Gooijer

**Author notes:** **To whom correspondence should be addressed**: M.C. de Gooijer; Division of Pharmacology, room H3.015; Plesmanlaan 121, 1066CX Amsterdam, The Netherlands.

## Abstract

**Purpose:** *BRAF*^V600^-mutated melanoma brain metastases (MBMs) are responsive to BRAF inhibitors, but responses are generally less durable than those of extracranial metastases. We here tested the hypothesis that the drug efflux transporters P-glycoprotein (P-gp; ABCB1) and breast cancer resistance protein (BCRP;ABCG2) expressed at the blood–brain barrier (BBB) offer MBMs protection from therapy.

**Methods:** We intracranially implanted A375 tumor cells in wild-type and *Abcb1a/b;Abcg2*^-/-^ mice. We characterized the tumor BBB, analyzed drug levels in plasma and brain lesions after oral vemurafenib administration and determined the efficacy against brain metastases and subcutaneous lesions.

**Results:** Although contrast-enhanced MRI demonstrated that the integrity of the BBB is disrupted in A375 MBMs, vemurafenib achieved greater antitumor efficacy against MBMs in *Abcb1a/b;Abcg2^-/-^* mice compared to wild-type mice. Concordantly, P-gp and BCRP are expressed in MBM-associated brain endothelium both in patients and in A375 xenografts and limited vemurafenib penetration into A375 MBMs. Confirming the BBB-specific context of this protection, vemurafenib was equally effective against subcutaneous A375 tumors in WT and *Abcb1a/b;Abcg2^-/-^* mice. Intriguingly, although initially responsive, A375 MBMs rapidly developed therapy resistance, even in *Abcb1a/b;Abcg2^-/-^* mice, and this was unrelated to pharmacokinetic or target inhibition issues. Rather, MBMs likely resorted to noncanonical growth signaling, as target inhibition of canonical MAPK pathway signaling components was maintained in resistant intracranial A375 tumors.

**Conclusions:** We demonstrate that BRAF^V600E^-driven MBMs are partly intrinsically protected from vemurafenib by the BBB. Intriguingly, MBMs can also rapidly acquire resistance *in situ*, likely by resorting to non-canonical growth signaling.

## 1. INTRODUCTION

Metastatic melanoma patients historically have a poor survival, mostly due to a lack of available effective chemotherapeutics (1). However, in the last decade significant advances have been made that considerably improved the outlook for metastatic melanoma patients. These advances were kick-started by the discovery of oncogenic *BRAF* mutations as an important driver in melanoma (2). The vast majority of *BRAF* mutations are in codon 600, substituting a valine for either a glutamic acid, lysine, arginine or aspartic acid, and result in constitutively active BRAF^V600E/K/R/D^ oncoproteins (3). BRAF^V600^-positive melanoma cells can therefore proliferate independently from external growth stimuli [2]. Importantly, BRAF^V600^ can be therapeutically targeted, and three drugs have now been approved for treatment of metastatic melanoma: vemurafenib (4), dabrafenib (5) and encorafenib (6). All three BRAF^V600^ inhibitors have generated striking clinical responses and significantly improved survival of metastatic melanoma patients (7–9). However, therapy resistance invariably occurs, in most cases due to selection and outgrowth of clones that carry additional mutations more downstream in the MAPK signaling pathway (10). Therefore, BRAF inhibitors are currently successfully combined with MEK inhibitors (vemurafenib and cobimetinib (11), dabrafenib and trametinib (12), encorafenib and binimetinib (6)), yielding further improved response rates and survival (13–15).

Despite all the recent success in treatment of metastatic melanoma, it is still unclear whether patients with melanoma brain metastases (MBMs) benefit similarly from BRAF^V600^ inhibitors as metastatic melanoma patients with extracranial metastases. In the clinical studies that led to the approval of BRAF^V600^ inhibitors for metastatic melanoma, MBM patients were excluded from study participation. Only recently, clinical trials focusing specifically on MBM patients have been set up, and the results from several phase II trials appear to suggest that BRAF^V600^ inhibitors also induce responses in MBMs (16,17). However, these responses were generally shorter than those achieved in extracranial metastases, suggesting that resistance occurs in MBMs even more rapidly than in extracranial metastases (17). The reason for this rapid resistance in MBMs in unclear, but could be related to the brain environment. Several preclinical studies have demonstrated that vemurafenib (18,19), dabrafenib (20) and encorafenib (21) exhibit very poor brain penetration in mice as a result of efficient efflux by P-glycoprotein (P-gp; ABCB1) and breast cancer resistance protein (BCRP; ABCG2) at the blood–brain barrier (BBB). These observations seem to be corroborated by a clinical study showing that the concentration of vemurafenib in cerebrospinal fluid (CSF) was less than 1% of the plasma concentration (22).

The BBB limits the brain penetration of many xenobiotics, including many anticancer agents (23), and can consequently impact the intracranial anticancer efficacy of small molecule drugs (24,25). Importantly, drug efflux transporters can restrict drug delivery and efficacy even when the BBB is considered ‘leaky’ (26). We therefore here investigate the impact of P-gp and BCRP on the efficacy of vemurafenib against MBMs in an intracranial mouse model of BRAF^V600E^-driven melanoma. In line with the clinical data, we find that MBMs can respond to vemurafenib, most likely as a result of compromised BBB integrity. However, vemurafenib achieved greater antitumor responses in *Abcb1a/b;Abcg2^-/-^* mice, indicating that P-gp and BCRP still play a protective role at the compromised BBB of MBMs. Supporting this hypothesis, we find that P-gp and BCRP are expressed in MBM-associated brain endothelium both in patients and orthotopic xenografts in mice. Importantly, vemurafenib efficacy could be improved by coadministration of the P-gp/BCRP inhibitor elacridar, offering a potential clinical strategy for increasing vemurafenib efficacy against MBMs. Intriguingly, we also observed much more rapid therapy resistance in the preclinical MBM model compared to previously published extracranial melanoma mouse models, analogous to clinical observations. We conclude that BRAF^V600^-positive MBMs are not only less responsive to vemurafenib because drug efflux transporters at the BBB limit drug penetration into the tumor, but also because they can rapidly acquire resistance during treatment. Therefore, P-gp/BCRP inhibitors might help to improve the clinical response of MBMs to vemurafenib by increasing its brain penetration, but pinpointing the mechanism behind the brain-specific acquired resistance will likely be necessary to produce durable responses.

## 2. METHODS

### 2.1 Cell culture and drugs

A375 cells (RRID: CVCL_0132) expressing firefly luciferase and mCherry (A375-FM) were cultured in minimum essential medium (MEM) supplemented with 10% fetal bovine serum (FBS), 1% penicillin/streptomycin, L-glutamine, nonessential amino acids, sodium pyruvate and MEM vitamins (all Life Technologies, Carlsbad, CA). The origin of the A375-FM cells was authenticated using STR analysis. Vemurafenib was purchased from LC Laboratories (Woburn, MA), vemurafenib-^13^C6 was obtained from the Slotervaart Hospital pharmacy and elacridar was generously provided by GlaxoSmithKline (Research Triangle Park, NC).

### 2.2 Animals

Mice were housed and handled according to institutional guidelines complying with Dutch and European legislation. All experiments with animals were approved by the animal experiment committee of the institute. The animals were athymic (nude) mice of a > 99% FVB background with wild-type (WT) or *Abcb1a/b;Abcg2^-/-^* genotype, between 8 and 12 weeks of age. The animals were kept in a temperature-controlled environment at 20.9 °C on a 12 hour light/dark cycle and received chow and acidified water *ad libitum*.

### 2.3 Drug formulations

A stock solution (25 mg/ml or 10 mg/ml) of vemurafenib was dissolved in dimethyl sulfoxide (DMSO) and Cremophor EL (1:1; both Sigma-Aldrich; St, Louis, MO). The working solution (2.5 mg/ml or 1 mg/ml) was freshly prepared prior to administration by diluting the stock solution with saline on the day of administration. Elacridar (5mg/ml) was formulated in DMSO:Cremophor EL:water (1:2:7) and prepared similarly.

### 2.4 Xenograft models and tumor growth monitoring

For subcutaneous xenograft models, 30 μL of cell suspension containing 3 × 10^6^ A375-FM cells were injected into both flanks of WT and *Abcb1a/b;Abcg2^-/-^* nude mice. Stereotactic intracranial injections of melanoma cells were performed as described previously (24). Briefly, WT and *Abcb1a/Abcg2*^-/-^ nude mice were injected intracranially with 2 μL of A375-FM cell suspension containing 1 × 10^5^ cells 2 mm lateral, 1 mm anterior and 3 mm ventral from the bregma. Tumor growth was measured by bioluminescence imaging (BLI) for intracranial tumors and by caliper for subcutaneous tumors. The volume of subcutaneous tumors was calculated in mm^3^ using the modified ellipsoid formula (volume = 0.5 x length x width^2^). Bioluminescence images were acquired following i.p. D-luciferin (150 mg/kg; Promega; Madison, WI) using an IVIS 200 or IVIS Spectrum system with Living Image software v4.5 (both PerkinElmer; Waltham, MA). Animals were stratified into treatment groups to achieve a similar mean bioluminescence reading within each cohort. The bioluminescence intensity of each individual animal at the day of start of treatment (day 0) was arbitrarily set at 100%. All subsequent measurements were recorded relative to this first measurement and converted to their log-values. Mean ± standard error (SE) values were calculated and plotted in graphs.

### 2.5 Magnetic resonance imaging

A BioSpec 70/20 USR (Bruker, Billerica, MA) system was used for magnetic resonance imaging (MRI), as described previously (24). The MRI sequence consisted of T2-weighted, T1-weighted pre-contrast and T1-weighted post-contrast imaging. Gadoterate meglumine (Dotarem^®^; Guerbet; Villepinte, France) diluted five-fold with saline was used as a contrasting agent and delivered via an intravenous cannula inserted in the tail vein. Mice were anesthetized using isoflurane (Pharmachemie B.V., Haarlem, the Netherlands) delivered via a customized mouse holder, and heart rate and breathing frequency were monitored throughout the entire procedure. Paravision software (v 6.0.1; Bruker) was used for image acquisition and Fiji (27) (v 1.49b) was used for image processing.

### 2.6 Pharmacokinetic studies

To establish vemurafenib plasma kinetics, tumor-free FVB WT and *Abcb1a/b;Abcg2^-/-^* nude mice received vemurafenib orally (p.o.) by gavage at indicated doses. Elacridar (100 mg/kg) was administrated p.o. by gavage four hours before vemurafenib. Blood was sampled from the tail vein at 15 min, 1 h, 2 h, 4 h, 8 h and 24 h after the administration. The plasma was obtained by centrifugation (5 min, 5,000 rpm, 4°C). Vemurafenib was extracted from plasma by diethyl ether liquid-liquid extraction. Extracts were dried using a Savant SpeedVac Concentrator (Thermo Fisher Scientific; Waltham, MA), reconstituted in MeCN:water (30:70) and subjected to liquid chromatography/tandem mass spectrometry (LC/MS-MS) analysis. Vemurafenib-^13^C6 was used as an internal standard.

To study its distribution in tumor-bearing mice, vemurafenib was administered p.o. to tumor-bearing mice for three days at a dose of 10 mg/kg or 25 mg/kg q.d., starting 14 days after intracranial tumor cell injection. One group of mice received 10 mg/kg vemurafenib 4 hours after administration of 100 mg/kg elacridar. Four hours after the third administration, blood was collected by heart puncture, whole brains were dissected and divided into four parts: ipsilateral hemisphere (tissue from the tumor-bearing hemisphere that was free of macroscopic tumor tissue), contralateral hemisphere (the tumor-free hemisphere), cerebellum and macroscopic tumor. In a follow-up experiment tissues and plasma were collected 4 hours after 5 and 10 consecutive daily p.o. administrations of vemurafenib. All tissues were weighed and subsequently homogenized using a FastPrep^®^-24 (MP-Bio-medicals, NY, USA) in 3 ml 1% (w/v) bovine serum albumin. All tissue samples were prepared for LC-MS/MS analysis as described above for plasma samples.

### 2.7 LC-MS/MS analysis

The LC-MS/MS system consisted of an API 3000 mass spectrometer (Sciex, Framingham, MA) coupled to an UltiMate 3000 LC System (Dionex, Sunnyvale, CA). Samples were separated using a ZORBAX Extend-C18 column (Agilent, Santa Clara, CA), preceded by a Securityguard C18 pre-column (Phenomenex, Utrecht, The Netherlands). Elution was done using a mixture of mobile phase A (0.1% formic acid in water (v/v)) and mobile phase B (methanol) in a 5 minute gradient from 20% to 95%B, followed by 95%B that was maintained for 3 min and then reequilibrated at 20%B. Multiple reaction monitoring parameters were 490.2/383.1 (vemurafenib) and 496.2/389.1 (vemurafenib-^13^C6). System control and data analysis was done using Analyst^®^ 1.6.2 software (AB Sciex; Foster City, CA).

### 2.8 Efficacy studies in xenograft models

For the subcutaneous tumor model, therapy was initiated two weeks after implantation, when the tumor volume exceeded 40 mm^3^. FVB WT and *Abcb1a/b;Abcg2^-/-^* nude mice (n = 8) received 25 mg/kg and 10 mg/kg of vemurafenib daily, respectively. Control mice (n = 8) received vehicle. Tumor development was assessed by caliper twice a week. For the intracranial tumor model, treatment was started about two weeks after intracranial injection of tumor cells, when full-blown tumors were present in all animals. WT and *Abcb1a/b;Abcg2^-/-^* nude mice received vehicle, 25 mg/kg vemurafenib, 10 mg/kg vemurafenib plus 100 mg/kg elacridar or 10 mg/kg vemurafenib once daily for 10 consecutive days or in a 5 days on/2 days off/5 days on schedule, as indicated in the panels of Figure 2. Tumor growth was monitored by BLI every four or five days. Mice were weighed daily weighed examined for abnormalities. The mice were humanely sacrificed based on bioluminescence imaging results or when weight loss exceeded 20% of the initial body weight.

**Figure 1.**
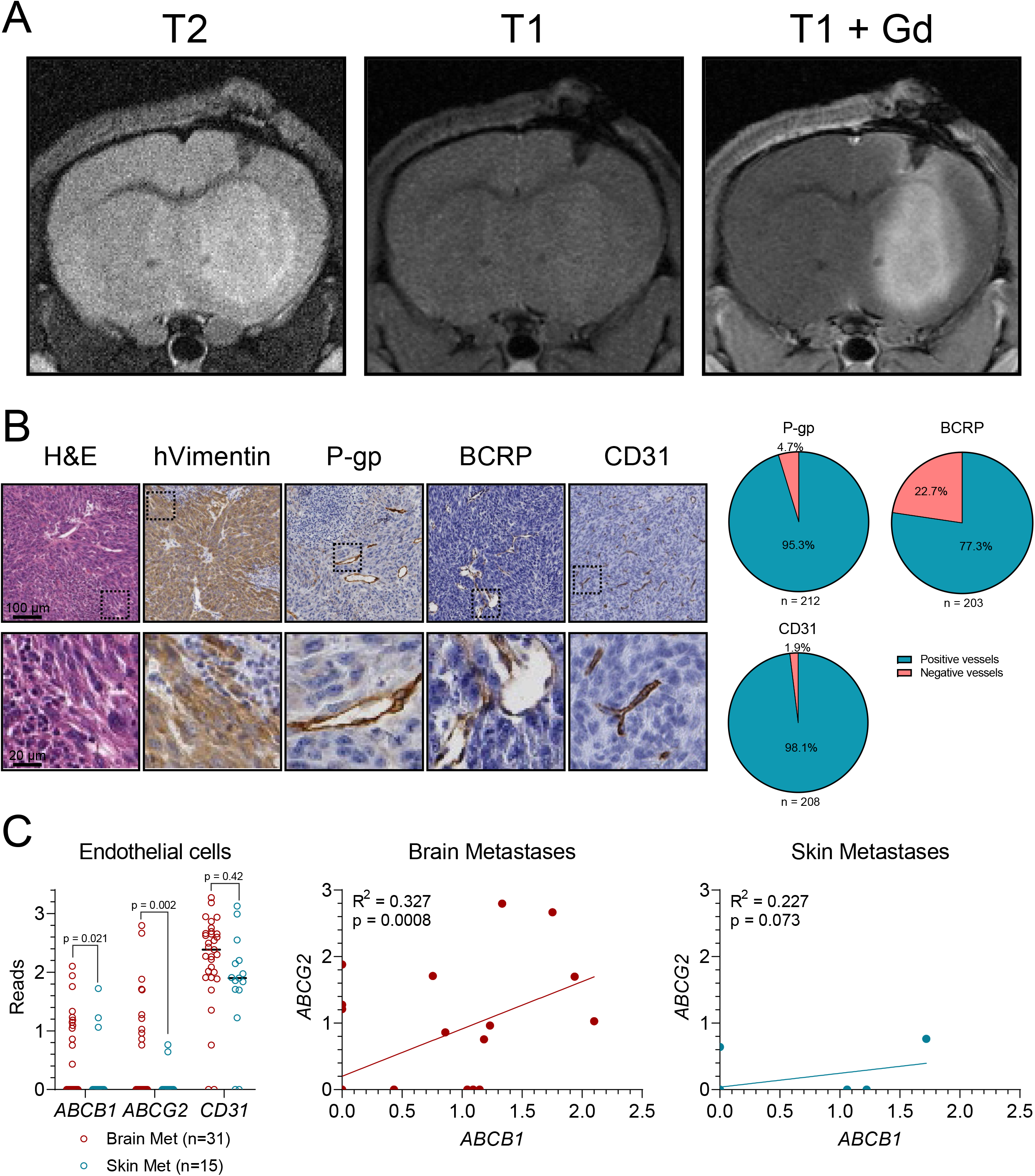
Characterization of the A375 melanoma brain metastasis model. (**A**) T2-weighted, T1-weighted pre-contrast and T1-weighted post-gadolinium (Gd) contrast magnetic resonance imaging of A375 tumors grafted in the brains of WT nude mice. (**B**) Histochemical staining with hematoxylin and eosin (H&E) and immunohistochemical staining of human vimentin (hVimentin), P-glycoprotein (P-gp), breast cancer resistance protein (BCRP) and CD31 of intracranial A375 melanoma tumors. Pie charts represent quantifications of positively and negatively stained vessels within A375 tumors. (**C**) Analysis of *ABCB1, ABCG2* and *CD31* gene expression by endothelial cells in brain and skin lesions from metastatic melanoma patients. Single cell RNAseq data was reported by Smalley et al. (28)

**Figure 2.**
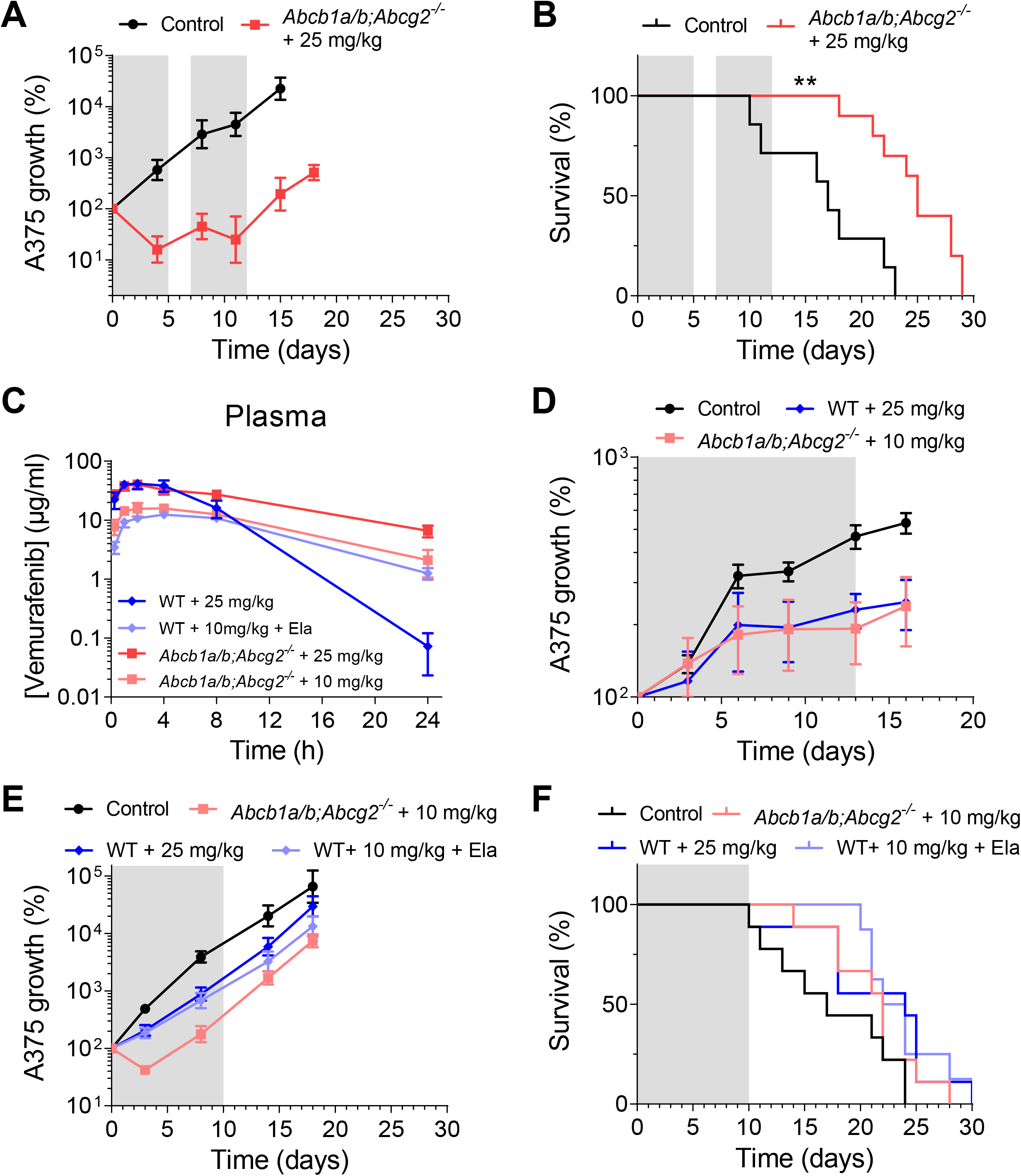
Intrinsic and acquired resistance of A375 tumors against vemurafenib *in vivo*. (**A**) Tumor growth and (**B**) survival of *Abcb1a/b;Abcg2^-/-^* mice bearing intracranial A375 melanoma tumors treated with two rounds of 25 mg/kg vemurafenib *q.d.x5d* or vehicle control. Treatment periods are shaded in grey. Data are represented as mean ± SD (n ≥ 7); ** *p* < 0.01. (**C**) Oral vemurafenib plasma concentration–time curves in WT mice receiving 25 mg/kg, WT mice receiving 10 mg/kg vemurafenib 4 hours after administration of the P-gp/BCRP inhibitor elacridar (Ela), *Abcb1a/b;Abcg2^-/-^* mice receiving 25 mg/kg vemurafenib and *Abcb1a/b;Abcg2^-/-^* mice receiving 10 mg/kg vemurafenib. Data are represented as mean ± SD (n ≥ 5). (**D**) Tumor growth of subcutaneous A375 tumors grafted in WT or *Abcb1a/b;Abcg2^-/-^* mice treated with various doses of vemurafenib administered *q.d*x13d or vehicle control. Treatment periods are shaded in grey. Data are represented as mean ± SD (n ≥ 7). (**E**) Tumor growth and (**F**) survival of WT and *Abcb1a/b;Abcg2^-/-^* mice bearing intracranial A375 melanoma tumors treated with 25 mg/kg, 10 mg/kg and 100 mg/kg elacridar (Ela) or 10 mg/kg vemurafenib *q.d*x10d or vehicle control. Treatment periods are shaded in grey. Data are represented as mean ± SD (n ≥ 8).

### 2.9 Histology, immunohistochemistry and image quantification

Mouse brains were fixed in 4% (v/v) formaldehyde. Alternatively, whole heads without skin were fixed in 4% (v/v) formaldehyde and 5% (v/v) glacial acetic acid and subsequently decalcified using a 6.5 % (v/v) formic acid solution for 3 days at 37 °C. After fixation, samples were paraffin embedded and sliced into 4 μm thick sections. Whole head slides were stained for hematoxylin and eosin (H&E), human vimentin (1:4,000; M0725; DakoCytomation; Glostrup, Denmark), P-gp (1:200; 13978; Cell Signaling Technology; Danvers, MA), BCRP (1:400; ab24115; Abcam; Cambridge, UK) and CD31 (1:200; ab28364; Abcam). Vessel positivity for P-gp, BCRP and CD31, as quantified in Figure 1B, was manually scored in at least 200 vessels per staining.

Brain slides were stained for H&E, PDGFRβ (1:50; 3169; Cell Signaling Technology), phospho-MET (1:150; 3077; Cell Signaling Technology), MET (1:100; AF527; R&D Systems; Minneapolis, MN), phospho-IGF1R (1:40,000; sc-101703; Santa Cruz Biotechnology; Dallas, TX), AXL (1:100; 8661; Cell Signaling Technology), NGFR (1:400; 8238; Cell Signaling Technology), phospho-EGFR (1:600; ab40815; Abcam), EGFR (1:200; ab52894; Abcam), BRAF^V600E^ (1:100; E19290; SpringBioscience; Pleasanton, CA), phospho-ERK1/2 (1:200; 4370; Cell Signaling Technology), phospho-AKT (1:8,000; 4060; Cell Signaling Technology, phospho-S6 (1:1,000; 2211; Cell Signaling Technology), phospho-4EBP1 (1:100; 2855; Cell Signaling Technology), 4EBP1 (1:1,200; 9644; Cell Signaling Technology), SOX10 (1:100; sc-17342; Santa Cruz Biotechnology), MITF (1:100; 284M-94; Cell Marque; Rocklin, CA) and Ki-67 (1:3,000; ab15580; Abcam).

### 2.10 Single cell RNAseq data analysis

Single cell RNAseq data from brain and skin lesions from metastatic melanoma patients as reported by Smalley *et al*. was accessed from http://iscva.moffitt.org (28).

### 2.11 Pharmacokinetic calculations and statistical analysis

Pharmacokinetic parameters were calculated with PKSolver (29). All comparisons involving more than two groups were analyzed by one-way ANOVA followed by Bonferroni *post hoc* tests. Differences in fractions of ABC transporter expressing cells between brain and skin lesions was compared using the Binomial test in which the skin fraction was considered as expected and the brain fraction was considered as observed. Correlations were determined using simple linear regression. Kaplan-Meier curves were drawn using GraphPad Prism v7 (GraphPad Software; La Jolla, CA) and statistically significant survival differences were determined using the log-rank test. Statistical significance was accepted in all tests when *p* < 0.05.

## 3. RESULTS

### 3.1 Characterization of the A375 melanoma brain metastasis model

To characterize the BBB integrity of the A375 MBM model, we subjected mice that were orthotopically injected with A375-FM cells to magnetic resonance imaging and (immuno)histochemical analysis. Similar to the clinical presentation of MBMs, intracranial A375 tumors were visible on T2-weighted and T1-weighted post-gadolinium contrast MRI sequences (Figure 1A). Enhancement on T1-weighted MR images after intravenous administration of a contrast agent indicates a reduction in BBB integrity. However, the BBB is not only a physical barrier but also a physiological barrier because of the expression of a range of efflux transporters, of which P-gp and BCRP are the most dominant. Immunohistochemical staining of these transporters in intracranial A375 tumors revealed that the majority of the vasculature in these tumors expresses P-gp and BCRP, as well as the endothelial cell marker CD31, suggesting that the physiological component of its BBB may still be functional (Figure 1B). Interestingly, intracranial A375 tumors were also characterized by large infiltrations of cells that resembled neutrophils, as apparent from their morphology and lack of staining for human vimentin. Melanomas are generally considered to be highly immunogenic, and widespread neutrophil infiltration could be a result of the immunogenicity of the A375 model.

To assess whether the orthotopic A375 model faithfully resembles ABC transporter expression at the BBB of MBMs in patients, we analyzed single cell RNAseq data from melanoma brain and skin metastases recently reported by Smalley *et al*. (28). Although these datasets unfortunately do not contain large numbers of endothelial cells, they suggest that P-gp/ABCB1 and BCRP/ABCG2 are expressed in MBM-associated endothelial cells, albeit heterogeneously (Figure 1C). Importantly, hardly any expression was found in endothelial cells from skin metastases, suggesting that these transporters do not have an impact on skin lesions. Finally, we could also observe that endothelial cells associated with brain metastases tended to co-express P-gp and BCRP to similar extents, while this correlation did not occur in the relatively few endothelial cells from skin metastases that express P-gp or BCRP. Together, the observations from the Smalley *et al*. dataset led us to conclude that the orthotopic A375 MBM model recapitulates the P-gp and BCRP expression found in MBM patients.

### 3.2 Vemurafenib has intrinsic antitumor potential against intracranial A375 tumors

The brain penetration of vemurafenib was previously reported to be significantly higher (between approximately 20- and 80-fold) in *Abcb1a/b;Abcg2^-/-^* compared to WT mice (18,19). We therefore first studied the efficacy of vemurafenib treatment against A375 tumors implanted in the brains of *Abcb1a/b;Abcg2^-/-^* mice, as we expected these mice to be the most pharmacologically favorable recipients to establish the intrinsic antitumor potential of vemurafenib against MBMs. Indeed, two cycles of 5 days of 25 mg/kg daily oral vemurafenib induced regression and subsequent tumor stasis of A375 tumors in these mice (Figure 2A). When the treatment was stopped, tumor growth started at a similar speed as untreated tumors, but a survival difference was already established (Figure 2B), indicating that vemurafenib is intrinsically potent against MBMs.

### 3.3 Dose adaptions between WT and *Abcb1a/b;Abcg2^-/-^* mice are needed to level the systemic exposure of vemurafenib between strains

The aim of this study was to assess the impact of P-gp and BCRP at the BBB on the intracranial efficacy of vemurafenib against MBMs by comparing WT and *Abcb1a/b;Abcg2^-/-^* mice. The systemic exposure and oral bioavailability of vemurafenib is known to be attenuated by P-gp and BCRP (18,19). This difference in systemic exposure may confound a fair comparison between the strains and a reduction of the dose in *Abcb1a/b;Abcg2^-/-^* mice was deemed necessary. The previous pharmacokinetic studies were conducted in tumor-free mice and used different formulations than the Cremophor-based formulation utilized in this study. Therefore, we first assessed the plasma exposure in tumor-free WT and *Abcb1a/b;Abcg2^-/-^* mice receiving vemurafenib in a Cremophor-based formulation. *Abcb1a/b;Abcg2^-/-^* mice received the same dose as WT mice (25 mg/kg) or a reduced dose (10 mg/kg). WT mice received the full dose (25 mg/kg) or the reduced dose (10 mg/kg) with concomitant administration of the P-gp/BCRP inhibitor elacridar (Figure 2C). We administered vemurafenib 4 hours after elacridar, as this is approximately the *t*_max_ of oral elacridar in mice. Similar to earlier studies, the plasma area under the curve (AUC) was significantly higher in *Abcb1a/b;Abcg2^-/-^* mice compared to WT mice receiving the same dose (Table 1). Notably, the terminal half-life of vemurafenib was considerably shorter in WT mice making accurate leveling between strains by dose adjustments difficult. Reducing the dose to 10 mg/kg in *Abcb1a/b;Abcg2^-/-^* mice resulted in a lower plasma AUC than WT at 25 mg/kg, but the trough levels were significantly higher. Co-administration of elacridar to WT mice yielded a vemurafenib plasma exposure similar to that in *Abcb1a/b;Abcg2^-/-^* mice, suggesting that elacridar efficiently inhibits systemic clearance mediated by P-gp and BCRP.

**Table 1.** Pharmacokinetic parameters of vemurafenib after oral administration of different doses to WT and *Abcb1a/b;Abcg2*^-/-^ FVB mice. AUC, area under the curve; *C*_max_, maximum concentration in plasma; *t*_max_, time to reach maximum plasma concentration; *t*_1/2_, elimination half-life; *CL/F*, apparent clearance after oral administration. Data are represented as mean ± SD (n ≥ 5); \**p* < 0.05, \*\**p* < 0.01, \*\*\*\**p* < 0.0001, compared to WT mice receiving the same vemurafenib dose.

| Parameter | Time (h) | WT<br>25mg/kg | *Abcb1a/b*;<br>*Abcg2*<sup>-/-</sup><br>25mg/kg | WT<br>10mg/kg<br>+ elacridar | *Abcb1a/b*;<br>*Abcg2*<sup>-/-</sup><br>10mg/kg |
| --- | --- | --- | --- | --- | --- |
| AUC<sub>plasma</sub> (μg/ml·h) | 0-4 | 147 ± 28 | 140 ± 16 | 38 ± 5.7 | 56 ± 7.2* |
| AUC<sub>plasma</sub> (μg/ml·h) | 0-24 | 390 ± 98 | 530 ± 68* | 180 ± 24 | 230 ± 35 |
| AUC<sub>plasma</sub> (μg/ml·h) | 0-∞ | 390 ± 99 | 610 ± 84** | 190 ± 23 | 250 ± 49 |
| C<sub>max</sub> (μg/ml) |  | 42 ± 7.7 | 41 ± 4.5 | 12 ± 1.7 | 17 ± 1.7* |
| t<sub>max</sub> (h) |  | 2.6 ± 1.3 | 1.9 ± 0.9 | 3.6 ± 0.9 | 2.8 ± 1.1 |
| t<sub>1/2</sub> (h) |  | 2.1 ± 0.2 | 8.5 ± 1.2**** | 5.8 ± 0.6 | 6.7 ± 1.4 |
| CL/F (L/kg·h) |  | 0.07 ± 0.017 | 0.04 ± 0.006** | 0.05 ± 0.006 | 0.04 ± 0.006 |

In order to assess if the dose leveling between the strains was appropriate, we treated subcutaneously grafted A375 tumors with 25 mg/kg vemurafenib in WT mice and 10 mg/kg vemurafenib in *Abcb1a/b;Abcg2^-/-^* mice for 13 consecutive days. Vemurafenib penetration into subcutaneous tumors similar in WT and *Abcb1a/b;Abcg2^-/-^* mice and as we found that vemurafenib was equally effective (Figure 2D), we selected these dose regimens for the efficacy study against intracranial A375 tumors.

### 3.4 P-gp and BCRP limit vemurafenib efficacy against intracranial tumors

We next grafted WT and *Abcb1a/b;Abcg2^-/-^* mice with intracranial A375 tumors, to study whether P-gp and BCRP at the BBB affect antitumor efficacy in an MBM model. We again treated WT mice with 25 mg/kg and used 10 mg/kg vemurafenib for *Abcb1a/b;Abcg2^-/-^* mice. We also added a group of WT mice receiving 10 mg/kg with concomitant elacridar. In this case, we now found that vemurafenib was more effective in *Abcb1a/b;Abcg2^-/-^* than in WT mice (Figure 2E). While vemurafenib only reduced A375 growth speed in WT mice, it induced tumor regression in *Abcb1a/b;Abcg2^-/-^* mice during the first three days of treatment. Notably, however, while still under therapy, regrowth occurred in these mice reaching a similar tumor growth speed as in untreated animals before the completion of treatment. As a result, survival was not significantly extended (Figure 2F). Pharmacological inhibition of P-gp and BCRP by elacridar was less efficacious, as the vemurafenib antitumor efficacy was greater in *Abcb1a/b;Abcg2^-/-^* mice receiving 10 mg/kg than in WT mice receiving elacridar and the same dose of vemurafenib (Figure 2E). Taken together, these data indicate that P-gp and BCRP at the BBB can diminish the efficacy of vemurafenib against MBMs.

### 3.5 P-gp and BCRP reduce vemurafenib penetration in MBMs

P-gp and BCRP limit the brain penetration of vemurafenib by virtue of their efflux function at the BBB (18,19). However, it is unknown if the penetration into MBMs is similarly affected, as these lesions display signs of a compromised BBB on contrast-enhanced MRI. We therefore measured the vemurafenib distribution in tumor-bearing WT and *Abcb1a/b;Abcg2^-/-^* mice after 3 daily administrations of vemurafenib. We collected brain, tumor and plasma samples at approximately the *t*_max_ of vemurafenib (4 hours after the last administration). The vemurafenib plasma concentration in WT mice receiving 25 mg/kg was about 1/4^th^ of the concentration observed in our previous pharmacokinetic experiment (Figure 2C and Figure 3A), whereas much smaller discrepancies were observed in *Abcb1a/b;Abcg2^-/-^* mice (2-fold) and WT mice that also received elacridar (no difference). Notably, these tumor-bearing mice in the later experiment received three administrations of vemurafenib and the tumor-free mice in the earlier experiment only one. Therefore, these data could suggest induction of P-gp and BCRP by repeated vemurafenib administration, resulting in increased clearance.

**Figure 3.**
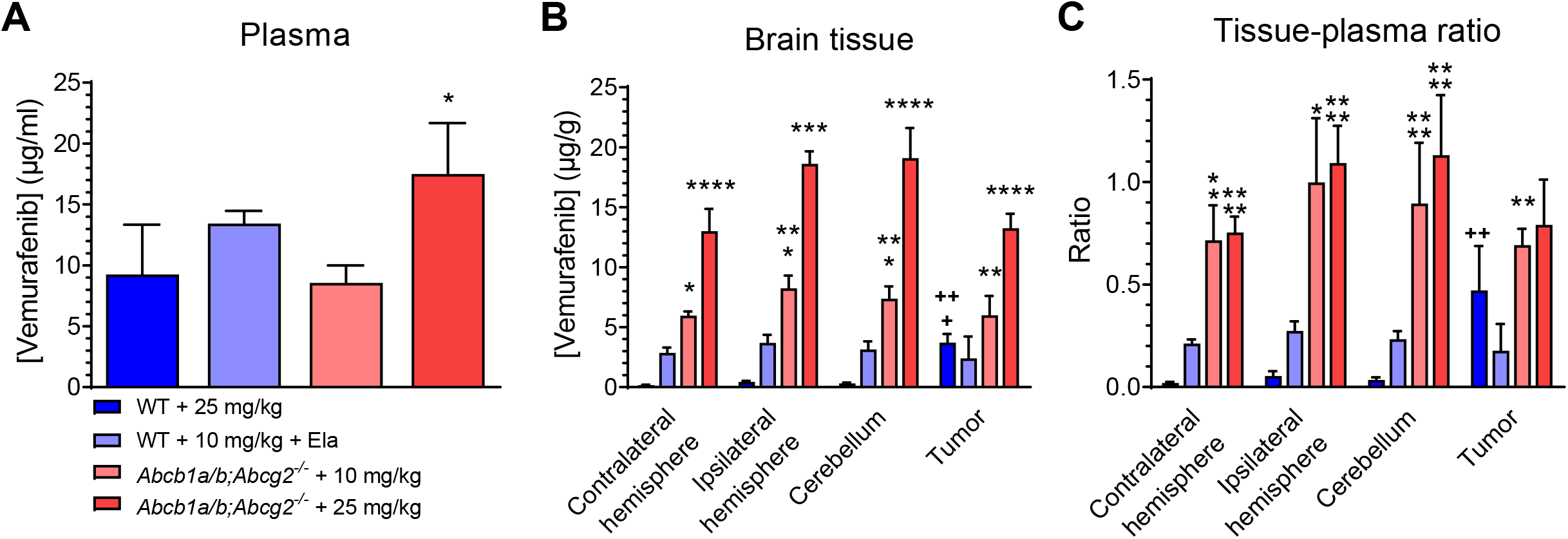
Vemurafenib concentrations in tumor and healthy brain of WT and *Abcb1a/b;Abcg2^-/-^* mice. (**A**) Plasma concentrations, (**B**) brain tissue concentrations and (**C**) tissue–plasma ratios in WT and *Abcb1a/b;Abcg2^-/-^* mice 4 hours after receiving oral vemurafenib at doses of 25 mg/kg, 10 mg/kg or 10 mg/kg 4 hours after receiving 100 mg/kg oral elacridar (Ela). The contralateral hemisphere represents the tumor-free hemisphere. The ipsilateral hemisphere is the hemisphere where the tumor was injected, from which all macroscopic tumor was removed. Data are represented as mean ± SD (n ≥ 3); \**p* < 0.05, \*\**p* < 0.01, \*\*\*\**p* < 0.0001, compared to WT mice receiving 25 mg/kg vemurafenib; ^++^ *p* < 0.01, ^+++^ *p* < 0.001, compared to the contralateral hemisphere level of the same group.

The apparent discrepancies in plasma concentration do not affect the results of the brain penetration as we always assess tissue–plasma ratio within each mouse. The vemurafenib concentrations in different brain regions differed greatly amongst all treatment groups (Figure 3B). As expected the highest concentrations were reached in *Abcb1a/b;Abcg2^-/-^* mice receiving 25 mg/kg vemurafenib. The concentrations were lower in *Abcb1a/b;Abcg2^-/-^* mice receiving 10 mg/kg vemurafenib, but this was only a result of the lower dose, as tissue–plasma ratios were similar between both dose levels in *Abcb1a/b;Abcg2^-/-^* mice (Figure 3C). In line with previous reports, the vemurafenib penetration in normal brain regions of WT mice was negligible. The tissue–plasma ratios were very close to the total blood volume of the murine brain (approximately 2%). Elacridar increased the vemurafenib concentration in healthy brain regions, but inhibition of P-gp and BCRP was incomplete, since the levels and brain-to-plasma ratios were significantly lower than in *Abcb1a/b;Abcg2^-/-^* mice. Vemurafenib penetrated into the tumor core in WT mice, but the levels were approximately half of those achieved in *Abcb1a/b;Abcg2^-/-^* mice. Elacridar was also not able to improve the penetration of vemurafenib into the tumor core to the same level as in *Abcb1a/b;Abcg2^-/-^* mice. These data show that P-gp and BCRP can still limit vemurafenib penetration into MBMs, even when the tumor lesion has compromised BBB integrity. These drug distribution data are in line with the observed intracranial antitumor efficacy (Figure 2A and 2E), as vemurafenib tumor concentrations were similar between WT mice receiving 25 mg/kg and WT mice receiving 10 mg/kg of vemurafenib with concomitant elacridar, but lower than in *Abcb1a/b;Abcg2^-/-^* mice receiving 10 mg/kg or 25 mg/kg vemurafenib.

### 3.6 Intracranial A375 tumors develop therapy resistance despite sufficient vemurafenib tumor penetration and target inhibition

As mentioned above, the A375 MBM model is responsive to vemurafenib, but developed therapy resistance after just a few days of treatment in *Abcb1a/b;Abcg2^-/-^* mice receiving 10mg/kg vemurafenib (Figure 2E). Since P-gp and BCRP are absent in these mice, we reasoned that P-gp/BCRP-unrelated pharmacokinetic phenomena may underlie the observed resistance. For instance, induction of vemurafenib pre-systemic metabolism, systemic clearance or efflux at the BBB by other ABC transporters might result in diminished vemurafenib brain concentrations after repeated administrations. However, the vemurafenib concentrations in various brain regions in tumor-bearing mice treated for a short (5 days) or long (10 days) period were not different in *Abcb1a/b;Abcg2^-/-^* mice and WT mice also receiving elacridar (Figure 4A-D). In fact, in contrast to the observed antitumor efficacy at 5 and 10 days of treatment (Figure 2E), the vemurafenib concentration in the tumor regions of these mice was even higher at the later time point (Figure 4E). Notably, we did find a considerably lower vemurafenib concentration in plasma in long-term treated WT mice compared to short-term treated WT mice (Figure 4A). Brain and tumor concentrations were also lower as a consequence of the lower plasma concentration, as the tissue–plasma ratios were unchanged over time (Figure 4F). Again, we found no reduction in vemurafenib plasma concentration in *Abcb1a/b;Abcg2^-/-^* mice or WT mice also receiving elacridar, indicating that the reduction in vemurafenib concentration over time in WT mice was mediated by P-gp and/or BCRP.

**Figure 4.**
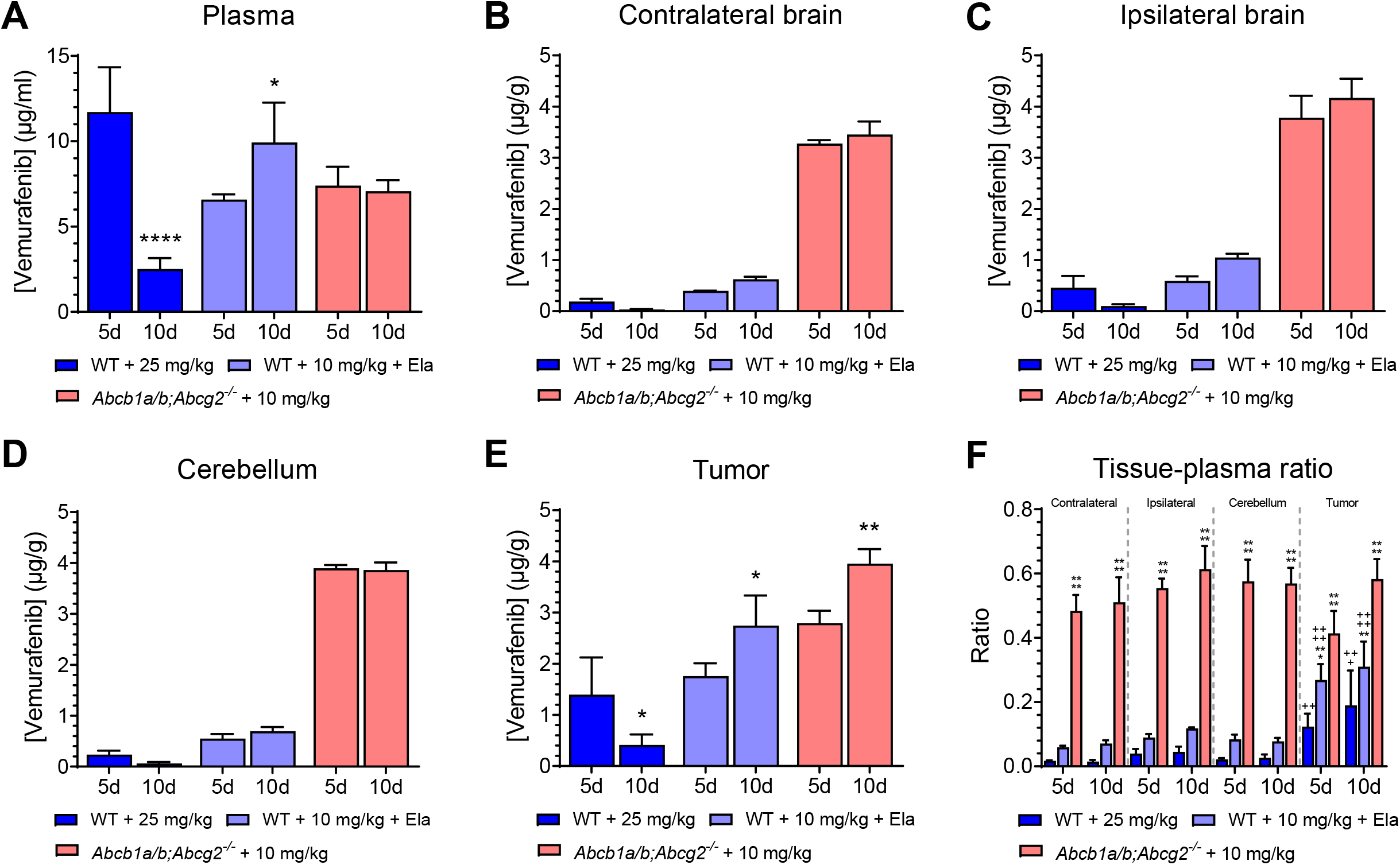
Vemurafenib concentrations in short-term and long-term treated intracranial A375 tumors. Vemurafenib concentrations in (**A**) plasma, (**B**) contralateral brain, (**C**) ipsilateral brain, (**D**) cerebellum and (**E**) tumor after short-term (5 days; 5d) and long-term treatment (10 days; 10d) of A375 melanomas brain metastases. Tumor-bearing WT or *Abcb1a/b;Abcg2^-/-^* mice orally received 25 mg/kg, 10 mg/kg or 10 mg/kg vemurafenib and 100 mg/kg oral elacridar (Ela). Data are represented as mean ± SD (n ≥ 3); \**p* < 0.05, \*\**p* < 0.01, \*\*\*\**p* < 0.0001, compared to short-term treated tumors of the same group. (**F**) Vemurafenib tissue–plasma ratios for different brain regions in 5 and 10 day-treated tumors. Data are represented as mean ± SD (n ≥ 3); ** *p* < 0.01, *** *p* < 0.001, **** *p* < 0.0001, compared to WT animals treated with 25 mg/kg vemurafenib; ^++^ *p* < 0.01, +++ *p* < 0.001, ^++++^ *p* < 0.0001, compared to the contralateral hemisphere region within the same treatment group.

Since the vemurafenib concentration in responsive short-term (5 days) treated tumors and resistant long-term (10 days) treated tumors was similar, we explored alternative ways by which intracranial A375 tumor may acquire resistance in a small pilot cohort we had available for immunohistochemical analysis (n=2 per group). Even though the cohort sample size was too small to robustly detect subtle differences in expression, we expected that we would be able to observe whether any substantial biological effects occurred. For instance, we observed canonical pathway inhibition in vemurafenib-resistant MBMs, as indicated by the profoundly reduced immunohistochemical staining of downstream BRAF^V600E^ targets phospho-S6 and phospho-4EBP1 (Figure 5). BRAF^V600E^,phospho-ERK and phospho-AKT were still low or unaffected, suggesting that resistance occurred via non-canonical growth signaling, as the proliferation marker Ki-67 was similarly unaffected. Upstream growth factor receptors are likely candidates for such a mechanism and have been demonstrated to mediate resistance to BRAF^V600^ inhibitors before (30–33). However, PDGFRβ, AXL, NGFR, MET and EGFR expression did not seem to be increased in resistant tumors and neither was signaling through phospho-IGF1R, phospho-MET or phospho-EGFR. In fact, PDGFRβ expression appeared to be diminished by vemurafenib treatment. Furthermore, we could not detect large differences in expression of transcription factors that have been implicated in acquired resistance mechanisms such as SOX10 (30) and MITF (32). Taken together, these findings suggest that rapid resistance in intracranial A375 tumors does not occur via pharmacological processes but through acquiring previously unreported non-canonical growth signaling.

**Figure 5.**
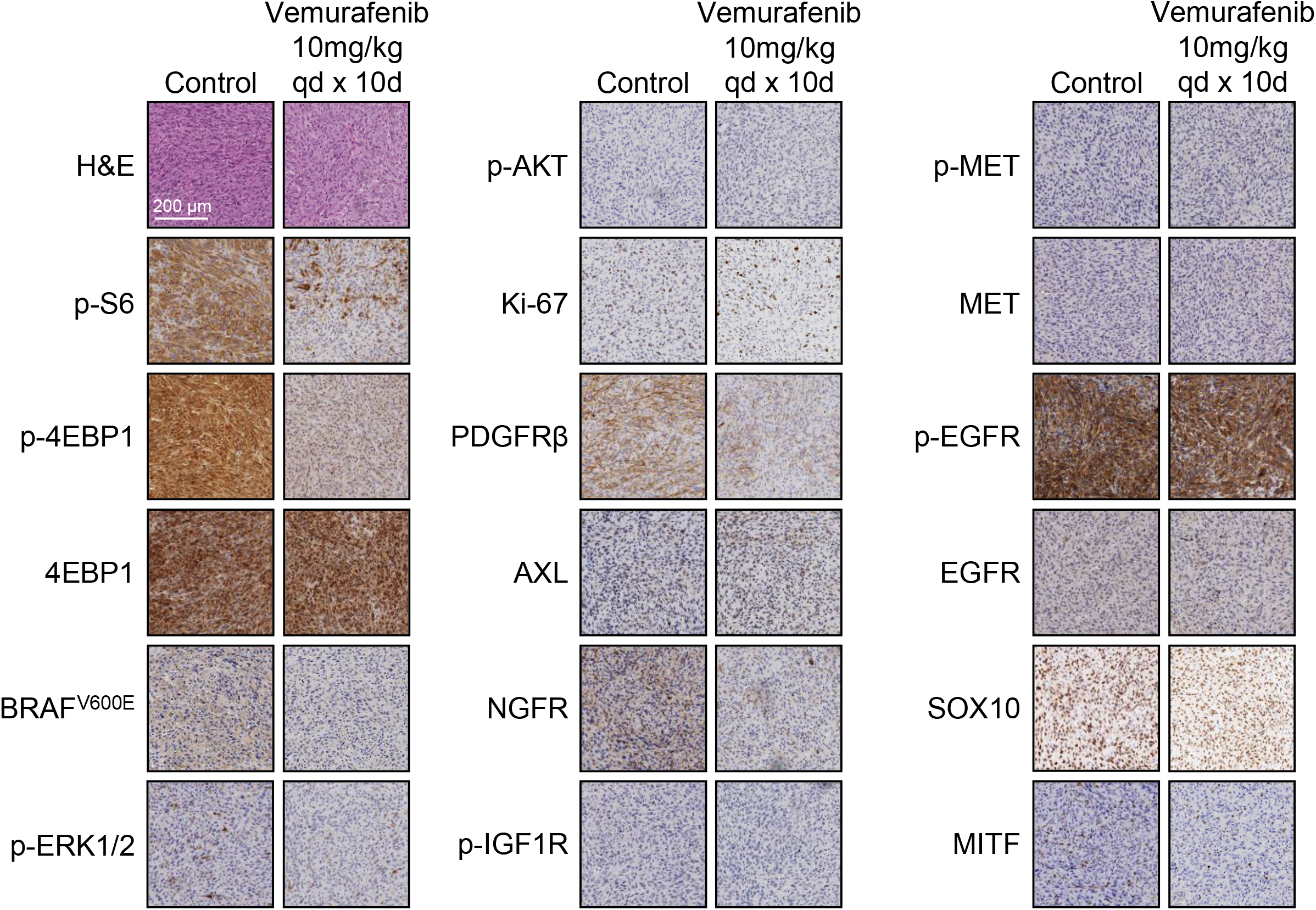
Immunohistochemical analysis of vemurafenib-resistant A375 melanoma brain metastases. Tumors growing in *Abcb1a/b;Abcg2^-/-^* mice treated orally with 10 mg/kg vemurafenib x10d (resistant stage) were stained for various markers and compared to untreated tumors. H&E, hematoxylin and eosin; p-S6, phospho-ribosomal protein S6; p-4EBP1, phospho-eukaryotic translation initiation factor 4E-binding protein 1; p-ERK, phospho-extracellular single-regulated kinase; p-AKT, phospho-AKT; Ki-67, proliferation marker protein Ki-67; PDGFRβ, platelet-derived growth factor receptor β; AXL, tyrosine-protein kinase receptor UFO; NGFR, neural growth factor receptor; p-MET, phospho-MET; p-IGF1R, phospho-insulin-like growth factor 1 receptor; p-EGFR, phospho epidermal growth factor receptor; SOX10, sex determining region Y-box 10; MITF, microphtalmia transcription factor.

## 4. DISCUSSION

The introduction of BRAF and MEK inhibitors has dramatically improved the survival of metastatic melanoma patients. However, clinical responses in melanoma brain metastases are less durable than those in extracranial metastases, suggesting MBMs may be intrinsically resistant to therapy (17). By using a preclinical mouse model, we here show that although BRAF^V600E^-positive MBMs cause a disruption of BBB integrity, P-gp and BCRP are expressed in the tumor blood vessels, thereby reducing the efficacy of vemurafenib by limiting its distribution into MBM lesions. Furthermore, by using *Abcb1a/b;Abcg2^-/-^* mice we found that BRAF^V600E^-positive MBMs are initially responsive to vemurafenib in the absence of P-gp and BCRP. However, they rapidly acquire resistance in the brains of these mice. This acquired resistance is not due to reduced levels of vemurafenib in the tumor, also not after repeated exposure. Therefore BRAF^V600E^-positive MBMs must acquire resistance to therapy by resorting to non-canonical proliferation signaling. However, no evidence was found that this occurs via previously described resistance mechanisms in extracranial melanomas (30–33).

The BBB limits the brain penetration and antitumor efficacy of treatment for primary brain tumors such as glioblastoma and diffuse intrinsic pontine glioma (34). However, its impact on the treatment of brain metastases is less well established (35). Brain metastases usually demonstrate contrast enhancement on T1-weighted MR imaging, indicating a loss of BBB integrity. Moreover, MBMs grow as relatively circumscribed lesions without much invasion of surrounding brain and remain in the vicinity of the vasculature (36,37). Consequently, MBM cells are rarely found outside of the contrast-enhanced brain regions where the BBB is intact. Therefore, clinical responses can be observed with poorly brain-penetrable drugs such as vemurafenib. These responses lead some to conclude that the relevance of the BBB is limited in brain metastases. Contrast enhancement on MR images indeed indicates a physical disruption of the BBB integrity, as tight junctions normally prevent paracellular diffusion of contrast agents. However, despite this loss of integrity, the drug efflux transporters P-gp and BCRP can still be functional in these brain lesions (26). In line with this hypothesis, we observed increased efficacy of vemurafenib against a BRAF^V600E^-positive MBM model that displays T1-weighted MRI contrast enhancement when these tumors were grafted in *Abcb1a/b;Abcg2^-/-^* mice and when vemurafenib was combined with the P-gp/BCRP inhibitor elacridar in WT mice. Hereby, we showed that even when BBB integrity is lost, brain penetration and antitumor efficacy of targeted agents that are substrates of P-gp and/or BCRP can still be limited.

The BBB may thus limit the efficacy of BRAF inhibitors against BRAF-mutated tumors residing in the brain. These do not only include brain metastases of melanoma (38) and non-small cell lung cancer (39), but also subsets of several different of primary adult (40) and pediatric (41) brain tumors. The expression of P-gp and BCRP in vessels of primary brain tumors is well-documented (34,42). Unfortunately, there are only two papers on P-gp or BCRP expression in blood vessels of brain metastatic lesions. Richtig *et al*. reported a general lack of P-gp expression in MBMs (43), whereas the blood vessels of various subtypes of breast cancer brain metastases were positive for BCRP (44). The results in human MBMs are not in line with our results in mice. This may be related to the size of the lesion, as stainings in human samples were all done on relatively large lesions that may depend more on angiogenesis. Notably, BCRP may be a more important drug efflux transporter in humans than in mice, since it is more abundantly expressed (45).

To maximize the potential of BRAF inhibitor therapy against intracranial malignancies, it is important to optimize its pharmakinetic and pharmacodynamic parameters. In that regard, vemurafenib does not appear to be the superior BRAF inhibitor. Pharmacokinetically, the brain–plasma ratios of oral vemurafenib in WT mice are around 0.02 (18,19), for encorafenib roughly 0.004 (21) and for dabrafenib approximately 0.1 (20). While a brain–plasma ratio of 0.1 for dabrafenib is still quite poor, it is clearly better than those of vemurafenib and encorafenib. Dabrafenib is also pharmacodynamically superior, as its IC_50_ against A375 cells is 4 nM (46). Encorafenib is similary potent against A375 cells (IC_50_ = 4 nM), but the IC_50_ of vemurafenib is approximately 100-fold higher at rougly 500 nM (47,48). As a consequence of the higher potency, plasma levels of dabrafenib given at therapeutic doses are about 20- to 50-fold lower (49). Nevertheless, these data suggest that dabrafenib may be the inhibitor of choice for treatment of BRAF-mutated intracranial tumors. This notion seems to be supported by clinical data. MBM patients receiving vemurafenib had a median overall survival of 4.3 months (50), compared to 7 months for dabrafenib treatment alone (16). To what extent this superior overall survival can be attributed to the higher intrinsic potency of dabrafenib and how much to its higher brain penetration is unclear, but both characteristics are likely to have contributed. In summary, the currently available data seems to suggest that dabrafenib-based treatment regimens have superior efficacy and that co-adminstration of P-gp/BCRP inhibitors such as elacridar may further enhance their efficacy.

Next to reduced sensitivity caused by the BBB, we observed a striking development of acquired resistance that occurred much more rapidly than is typically reported for extracranial tumor models (51). Interestingly, these data seem to be in line with observations in metastastatic melanoma patients. In a phase II study investigating dabrafenib and trametinib combination therapy in metastatic melonoma patients with brain metastases, similar intracranial and extracranial response rates (approximately 50%) were observed (17). However, the duration of response was considerably shorter for intracranial metastases (6.5 months) than for extracranial metastases (10.2 months). The reason why MBMs acquire therapy resistance more rapidly is not yet understood. Several resistance mechanisms to BRAF inhibitors have been described to date (52). Notable mechanisms include increased EGFR signaling (30), increased PDGFRβ signaling (31) and a low MITF/AXL ratio (32). The previously reported effect sizes of these mechanisms are quite striking and since we could not detect any major changes in our pilot cohort of resistant intracranial A375 tumors these mechanisms appear not be implicated in the resistance we observed (Figure 5). A more recently discovered resistance mechanism is the acquisition of a secondary *BRAF* mutation resulting in a BRAF^V600E/L514V^ oncoprotein (53), but this is unlikely to occur in our A375 MBM model as this mutation would lead to increased canonical MAPK pathway signaling, which we did not observe. Microenvironment-related resistance mechanisms exerted by reactive astrocytes have also been proposed (37). For instance, factors secreted by astrocytes have been demonstrated to increase AKT signaling in melanoma cells *in vitro* (54). This specific mechanism is unlikely to have occurred in our study, as we did not observed increased p-AKT levels in resistant tumors (Figure 5). However, it does indicate that the microenvironment can contribute to acquired therapy resistance. Indeed, a potential role for the MBM microenvironment may also help to explain the observed differential clinical responses of intracranial and extracranial metastases (17).

Taken together, this study demonstrates that BRAF^V600E^-positive MBMs are not only less sensitive to vemurafenib because they are still partially protected by expression of P-gp and BCRP in the disrupted BBB, but they can also rapidly acquire resistance likely dependent on the unique microenvironment of the brain. Adding a P-gp/BCRP inhibitor to BRAF inhibitor therapy may therefore improve survival by overcoming intrinsic resistance of MBMs.. However, understanding the mechanism behind the apparent brain-specific acquired resistance will likely be necessary to induce long-term responses.

## Abbreviations used

AUC: area under the curve;
BBB: blood–brain barrier;
BCRP: breast cancer resistance protein;
CSF: cerebrospinal fluid;
MBM: melanoma brain metastasis;
MRI: magnetic resonance imaging;
P-gp: P-glycoprotein;
WT: wild-type

## Acknowledgements

We thank Hans Meel and Piotr Waranecki (Princess Maxima Center) for STR analysis of the A375-FM cell line.

